# Coxmos: Interpretable survival models for high-dimensional and multi-omic data

**DOI:** 10.1101/2025.10.01.679727

**Authors:** Pedro Salguero, Anabel Buendía-Galera, Sonia Tarazona

**Affiliations:** Department of Applied Statistics, Operations Research and Quality, Universitat Politècnica de València, Valencia, 46022, Spain

**Keywords:** survival, high-dimensional, multi-omics, variable selection, PLS

## Abstract

**Background:** Survival analysis in high-dimensional (HD) and multi-block (MB) settings, such as omic and multi-omic studies, poses major methodological challenges due to multicollinearity, low events-per-variable ratios, and limited model interpretability. While Cox Proportional Hazards models remain widely used, their applicability is restricted in HD scenarios. Machine learning approaches can handle such complexity but often lack interpretability. PLS-based survival models offer an attractive compromise, yet existing implementations provide limited support for model optimization, evaluation, and interpretation.

**Results:** To overcome these limitations, we introduce Coxmos, a CRAN R package that integrates adapted Cox regression models with HD-adapted variable selection, and optimized PLS-based approaches tailored for HD and MB data. Coxmos also provides validation, comparison, interpretation, and visualization tools. Benchmarking on clinical, single-omic, and multi-omic datasets demonstrated that Coxmos outperformed other state-of-the-art machine learning methods, while enhancing interpretability. With an ovarian cancer case study, we illustrated how Coxmos facilitates model selection, validation, and biological interpretation in a complex multi-omic survival analysis.

**Conclusions:** Coxmos provides a robust, flexible, and interpretable solution for survival analysis in HD and MB data. By combining optimized PLS-based methodologies with rigorous evaluation and rich visualization tools, Coxmos enables reliable survival prediction and facilitates the identification of relevant risk and protective factors in complex biomedical studies. The package fills an important methodological gap and represents a valuable resource for reproducible and interpretable survival modeling in the big data scenario.

**Availability:** Coxmos is available at CRAN repository: https://cran.r-project.org/web/packages/Coxmos.

## 1 Introduction

Survival analysis is a fundamental statistical approach in medicine and other fields to model the time until a specific event occurs (such as death or disease onset) as a function of other covariates (age, gender, treatment, etc.), while accounting for censored observations. The *Cox Proportional Hazards (PH) model* [[Cox, 1972]] remains one of the most widely used methodologies. This semiparametric model expresses the hazard at time *t* for an individual with covariate vector **x** as in Equation 1.

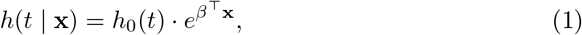

where *h*_0_(*t*) is the baseline hazard function and *β* the vector of coefficients associated to the covariates. The Cox model assumes that covariate effects are constant over time (PH assumption), requires that the number of observations *n* exceeds the number of predictors *p*, and cannot properly handle multicollinearity.

Besides survival prediction, an essential aim of survival models is to identify the most influential predictors of survival, a task that becomes increasingly complex as the number of predictors grows [Witten and Tibshirani, 2010]. This is the case of high-dimensional (HD) datasets, including omic data, where the number of variables *p* exceeds the number of observations *n*, multicollinearity is frequent, and the number of events per variable (EPV) —that determines the statistical power of the models— is too low [Concato et al., 1995, Peduzzi et al., 1995, Vittinghoff and McCulloch, 2007]. To address these limitations, variable selection techniques such as stepwise procedures or shrinkage methods like Elastic Net have been proposed [Guyon and Elisseeff, 2003].

Machine learning (ML) approaches are another alternative to model survival in HD settings [Spooner et al., 2020]. Techniques such as Support Vector Machines (SVM), Boosted Models, Random Forests, or Neural Networks have been adapted to handle right-censored data [Shivaswamy et al., 2007, Chen and Guestrin, 2016, Ishwaran et al., 2008, Faraggi and Simon, 1995, Hothorn, 2006, Knottenbelt et al., 2025]. These models often achieve high accuracy, and some incorporate feature selection strategies; however, their interpretability remains limited.

Conversely, dimensionality reduction techniques such as Partial Least Squares Regression (PLS) [Wold, 1966, Sjöström et al., 1983] provide a more interpretable alternative to deal with HD data. PLS models rely on a small number of latent variables that capture most of the variability in the original dataset and benefit from multi-collinearity. Various adaptations of PLS for survival analysis have been developed. [Nguyen and Rocke, 2002] applied the PLS model on survival time without considering censoring and used the extracted PLS components as predictors in a Cox PH model. [Park et al., 2002] reformulated the PLS algorithm to introduce Poisson regression to handle censoring. [Li and Gui, 2004] integrated Cox models within the PLS algorithm, thus considering censored observations. [Bastien, 2008] proposed the usage of deviance residuals from a null Cox model as the response variable in the PLS model, and this strategy was implemented in the *plsRcox* R package [Bastien et al., 2015], which also includes sparse (variable selection) and kernel-based (nonlinear relationship) versions. However, this package lacks tools for optimizing, evaluating and interpreting PLS-based survival models.

An even more complex scenario arises for multi-block (MB) datasets (e.g. from multi-omic experiments), where the multicollinearity issue increases and complicates the identification of relevant survival predictors. MB Random Forest [Hornung and Wright, 2019], IPF Lasso [Boulesteix et al., 2017] or Priority Lasso [Klau et al., 2018] are interesting options for MB datasets, while generally providing limited insight into the relative importance of individual omic layers. While MB-PLS regression models have been widely used for MB data [Westerhuis et al., 1998], to our knowledge, they have not yet been adapted for survival analysis.

Despite the variety of methodologies developed for survival analysis on HD or MB data, several challenges remain to be addressed, including overfitting, model interpretability, and the lack of robust tools for model evaluation and comparison. Solving these issues requires integrating efficient cross-validation procedures, predictive performance metrics, and advanced visualization tools to support model selection and results interpretation. To overcome these limitations, we have developed Coxmos, an R package providing a unified and flexible framework for survival analysis in HD and MB settings. Coxmos implements four modeling strategies for HD data: the Cox proportional hazards model with HD-adapted stepwise or Elastic Net variable selection, and three PLS-based approaches: PLS-ICOX, PLS-DRCOX, and PLSDACOX. It also extends the PLS approaches for survival modeling in MB studies. The package includes automated hyperparameter optimization via cross-validation, variable selection strategies, and different performance metrics for model optimization, evaluation and comparison.

## 2 Methods

### 2.1 PLS-based survival methodologies

Partial Least Squares (PLS) regression [Wold, 1966] models the relationship between a predictor matrix **X** and a response matrix **Y** through the linear model in Equation 2.

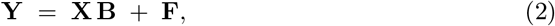

where **B** denotes the matrix of regression coefficients and **F** the residual matrix.

The strategy behind PLS is creating latent variables or components, as linear combinations of the original variables in **X** and **Y**, that maximize the covariance Cov(**t, u**), where **t** and **u** are the scores, that is, the coordinates of observations in the new latent space for **X** and **Y**, respectively, and represent the components. The **X** and **Y** matrices can be decomposed as in Equations 3 and 4:

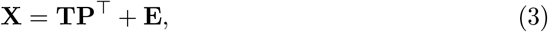

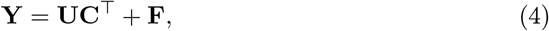

where **T** and **U** are the score matrices, **P** and **C** are the loading matrices (contributions of each variable to each component), and **E** and **F** are the residual matrices.

The NIPALS iterative procedure [Höskuldsson, 1988] can be used to obtain the PLS components, as follows:

1. Select a random column from **Y**, denoted as **u**.
2. Calculate the weight vector, **w** that maximizes the correlation between **X** and the scores of 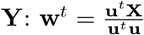.
3. Calculate **X**-component 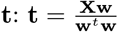.
4. Calculate **Y**-loading vector 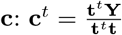.
5. Calculate **Y**-component 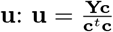.
6. Calculate **X**-loading vector 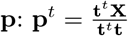.
7. Compute the residual matrices **E** = **X** − **tp**^*t*^ and **F** = **Y** − **tc**^*t*^.

For estimating a new PLS component *a* (*a* = 1, …, *A*), matrices **X** and **Y** from Step 2 must be replaced with the residuals **E** and **F**, respectively and proceed with the calculation of the corresponding loadings, scores or weights. This process continues until all components are computed. The number of components *A* to be retained should be optimized via cross-validation before computing the regression coefficients (Equation 5).

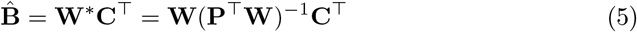

In survival analysis, PLS models must be adapted to handle both censoring information and survival times. Consequently, various options have been proposed, which also incorporate variable selection strategies to identify risk or protective factors in survival. The next sections describe the sparse PLS-based survival methods proposed and benchmarked in this work for the HD and MB scenarios.

#### 2.1.1 PLS-based survival models for HD data

##### sPLS-ICOX

The *sparse PLS individual Cox* (sPLS-ICOX) method in Coxmos was adapted from [Bastien et al., 2015]. In this approach, step 2 in NIPALS algorithm mentioned above is replaced with a univariate Cox regression model for each predictor *j*. Therefore, the weights are computed as 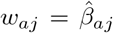, where 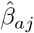 is the estimated Cox regression coefficient of predictor *j* in component *a*. The retained *A* PLS components are the predictors in the final multivariate Cox model. The score and loading vectors for **X** (**t** and **p**) are obtained following the expressions in Steps 3 and 6 of the original PLS algorithm, and the residual matrix **E** is computed or updated as in Step 7. This iterative process continues until the complete score matrix **T** is fully constructed. Steps 1, 4, and 5 are ignored in this approach.

In this work, we complement this methodology with a sparse approach to select the most relevant predictors. For that, we set the weight *w*_*aj*_ = 0 when the associated 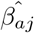 is not significant for a given significance threshold *α*, which is optimized by cross-validation (CV) strategies together with the number of PLS components *A* to be retained. Finally, these *A* components (**t**_**a**_, *a* = 1, …, *A*) are used as predictors in a multivariate Cox model, which returns the significant components for survival prediction.

##### sPLS-DRCOX

The *sparse PLS Deviance Residual Cox* (sPLS-DRCOX) method we present in this work is a modification of the splsRcox algorithm described in [Bastien, 2008]. In this algorithm, the PLS response variable is the vector of deviance residuals computed from a null Cox model, with no predictors. Prior to computing deviance residuals, Martingale residuals for each individual 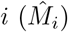 are obtained (Equation 6).

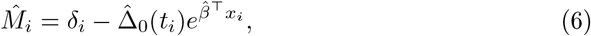

where *δ*_*i*_ is the event indicator and 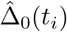 the estimated cumulative hazard function at time *t*_*i*_.

To circumvent the potential skewness attributed to Martingale residuals [Bastien et al., 2015], deviance residuals are computed (Equation 7).

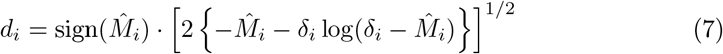

After fitting a PLS model on these deviance residuals, the resulting scores **t**_**a**_ (*a* = 1, …, *A*) are used as predictors in a final Cox model that provides the most significant components associated with survival.

In our sPLS-DRCOX method, we propose an alternative variable selection strategy, inspired by the Lasso variable selection in the *mixOmics* R package [Rohart et al., 2017]. The mixOmics sPLS method tries different combinations of the number of variables to retain for each PLS component *k*_*a*_ (*a* = 1, …, *A*) and chooses the one with the best performance metric. While a standard PLS analysis evaluates as many models as the number of components *A*, the mixOmics sPLS fits *A* models for each potential vector 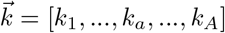. If *v* is the number of different values to be tried for each *k*_*a*_, the total number of models is 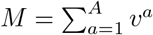. For omic datasets, with tens of thousands of variables, this strategy quickly becomes computationally prohibitive. A common solution is to drastically reduce the value of *v*, which often comes at the expense of excluding models that could provide better predictions.

To overcome this limitation, we designed a novel heuristic method that yields a near-optimal solution for 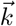 at a reasonable computational time. Our heuristic algorithm efficiently searches the solution space, instead of exhaustively testing all possible 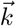. First, a small set of *C* evenly spaced numerical values (cutpoints) is selected within a specified range. This subset is used to fit initial models and evaluate their performance. If a particular configuration 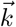 achieves the best performance, it is selected.

In the event of a tie, the configuration with the fewest variables is selected. Next, an iterative process explores the midpoints between this best configuration and its nearest neighbors until no further improvements are detected or the improvement is below a predefined threshold. This strategy reduces the number of computed models to *M* = *A* × (*C* + 2*I*), *I* is the number of additional iterations performed during the optimization. The maximum number of iterations, *I*, can be estimated as *I* ≈ log_2_(*R*), where *R* is the range of the numbers of variables to be tried, and could be even lower when no improvement is detected. This strategy drastically reduces the computational time to obtain a good solution for the variable selection problem.

##### sPLS-DACOX

The *sPLS-Discriminant Analysis* (sPLS-DACOX) strategy we propose in this work is based on the classification version of Partial Least Squares (PLS) models (PLS-DA). Our goal in this case is to define PLS components that maximize the separation between subjects who experienced the event and those who were censored. The time-to-event information is therefore not used in the PLS-DA model, but considered in a final Cox model where these PLS components act as predictors.

Our sPLS-DACOX method incorporates the same heuristic variable selection strategy described above for sPLS-DRCOX.

#### 2.1.2 PLS-based survival models for MB data

We present several survival methods for MB datasets. The first set of methods (*single-block*) analyzes each data block independently and combines the results in a final Cox model, while the second set of models (*multi-block*) jointly analyzes all data blocks.

##### Single-block sPLS methods for MB survival data

In this approach, each data block is first analyzed independently using any of the sPLS-based models previously described for HD data. Consequently, for each data block **X**_**b**_ (*b* = 1, …, *B*), we obtain *A*_*b*_ PLS components: 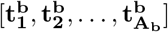. These 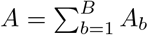 block components are then used as predictors in a multivariate Cox survival model.

We propose two variants of this method: SB and iSB. In the SB option, the number of components *A*_*b*_ and variable selection penalty are optimized via cross-validation; however, their values remain the same across all blocks. The number of the total components *A* is restricted by the EPV rule: 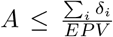, where *δ*_*i*_ is the censoring variable. In contrast, the iterative SB (iSB) approach performs cross-validation independently for each block, allowing for different numbers of components and penalties per block. In this case, each block can handle a maximum of *A*_*max*_ components, where 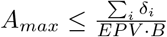.

##### Multi-block sPLS methods

The MB methods we present in this work simultaneously integrate all blocks within the same sPLS-based survival model [Tenenhaus and Tenenhaus, 2011, Tenenhaus et al., 2014]. Our MB-sPLS-DRCOX and MB-sPLS-DACOX methods are based on the MB PLS model implemented in the mixOmics R package [Rohart et al., 2017].

These methods rely on a block-correlation matrix that specifies the degree of relationship among blocks, which can either be defined according to prior knowledge or empirically estimated. We estimated them by computing the correlation between the first component from pairwise PLS models across all blocks.

### 2.2 Overview of Coxmos package

All the PLS-based survival methods described in the previous section were implemented in the Coxmos R package, which also includes functionalities for data exploration and preprocessing, assumption validation, performance evaluation and comparison, model interpretation, and prediction (Fig. 1). We also incorporated model optimization via cross-validation and advanced visualization capabilities to enhance model robustness and interpretability. All these functionalities are briefly described in the next sections.

**Figure 1:**
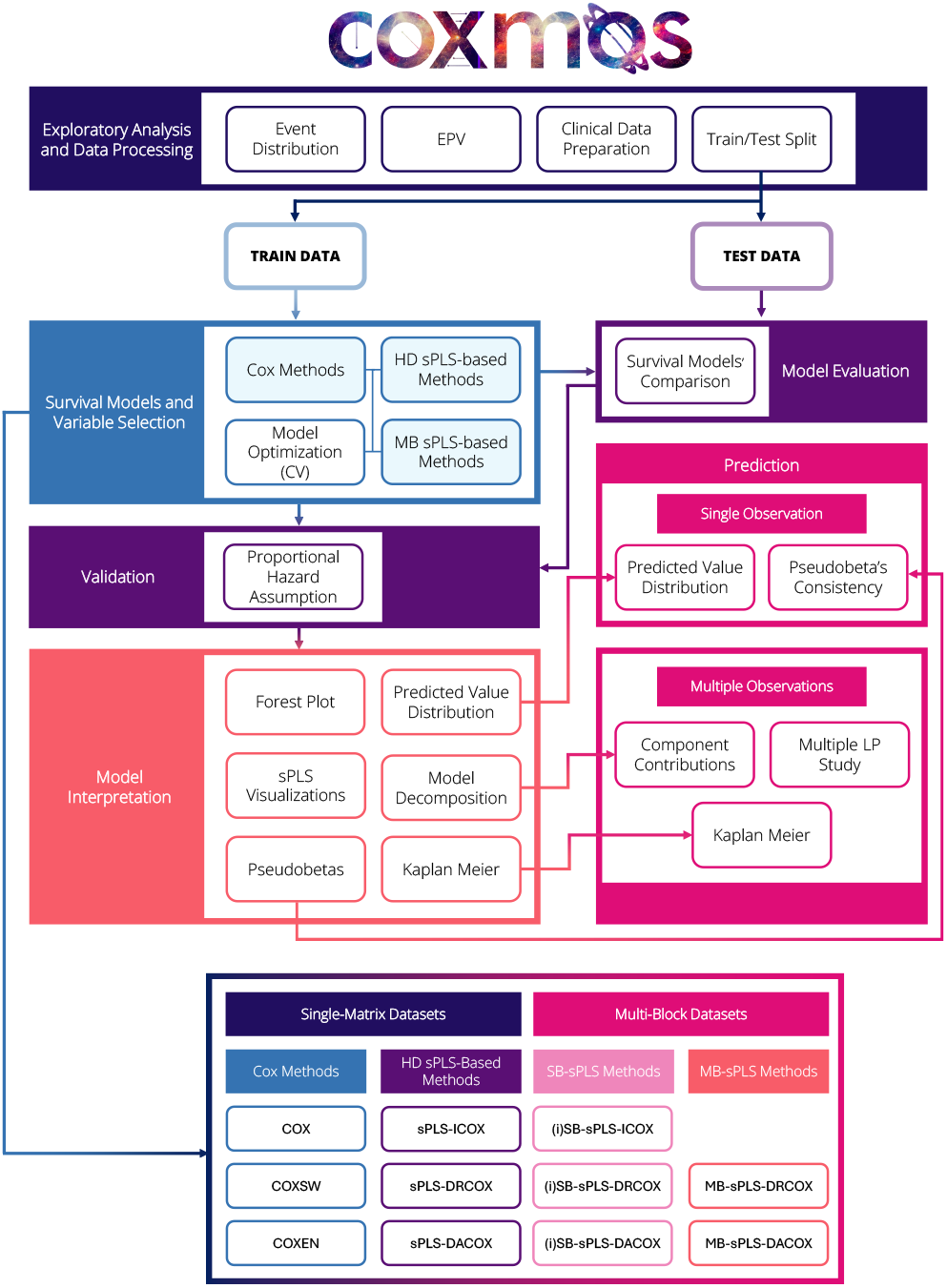
Overview of Coxmos package functionalities for survival analysis on high-dimensional (HD) and multi-block (MB) datasets.

#### 2.2.1 Data exploration and preparation

Before conducting survival analysis, Coxmos generates visualizations to assess data quality, showing the distribution of events and censored observations within a specified time range. As the stability and statistical power of the model depend on the EPV ratio and a value lower than 10 increases the risk of overfitting and unstable estimates, the package also reports the EPV for the analysed dataset.

Finally, prior to model fitting, Coxmos prepares the data by transforming categorical variables into dummy variables and creates a dataset partition into training and test subsets for proper validation that preserves the distribution of event/censored observations.

#### 2.2.2 Survival models in Coxmos

Coxmos implements all the PLS-based survival models described in the previous section: methods for HD datasets (sPLS-ICOX, sPLS-DRCOX and sPLS-DACOX) and models for MB datasets ((i)SB-sPLS-ICOX, (i)SB-sPLS-DRCOX and (i)SB-sPLS-DACOX, MB-sPLS-DRCOX and MB-sPLS-DACOX). In addition, Coxmos also incorporates the Cox PH model with two variable selection strategies adapted for HD data: stepwise and ElasticNet.

For all models, hyperparameters (penalty factors, number of PLS components, number of variables to retain per component, etc.) are optimized by applying a cross-validation procedure. Moreover, for the sake of transparency and reusability, the package returns all computed matrices (scores, residuals, loadings and weights).

##### Stepwise variable selection (CoxSW)

The stepwise (SW) method [Efroymson, 1960] —based on iteratively adding or removing variables— cannot always be used in HD scenarios since existing algorithms initialize from a full model with all the covariates, which cannot be fitted when *p* ≥ *n*. In addition, the SW algorithm may compromise the survival predictive performance when the EPV criterion is not met [Flom and Cassell, 2007, Thompson, 1995]. Therefore, we modified the SW proceduremfrom the *My*.*stepwise* R package [Company, 2017]. First, univariate Cox models are fitted for each covariate, rather than a full model. For the backward option, the initial model includes the most significant variables, as determined by their univariate p-values, while fulfilling the EPV constraint imposed by the user. The forward option adds variables with the most significant univariate p-values until the EPV constraint is no longer met. Finally, while *My*.*Stepwise* only relies on ANOVA tests to iteratively compare models; we combined ANOVA with Akaike’s Information Criterion (AIC) [Akaike, 1973] to select variables that contribute to a measurable improvement in model performance. This modification enhances algorithmic convergence, ensuring a more robust and consistent selection process.

##### ElasticNet strategy (CoxEN)

The ElasticNet (EN) regularization method [Simon et al., 2011] combines Lasso and Ridge approaches into the penalization term in Equation 8:

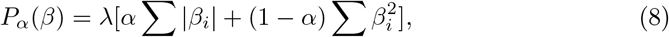

where *λ* is the regularization parameter and *α* is a value between 0 and 1 which gives the weights for Lasso and Ridge penalizations. We adapted the EN method in the *glmnet* R package [Simon et al., 2011] to include the EPV restriction and a custom cross-validation (CV) to also optimize the *α* parameter, since only *λ* optimization is performed in *glmnet*.

#### 2.2.3 Model optimization and evaluation

Coxmos applies a repeated k-fold CV on the training data to optimize model hyperparameters, where the number of folds and repetitions can be tuned. The model optimization is based on the performance metric of choice, or even on a weighted combination of several metrics. The performance metrics included in Coxmos are the Akaike Information Criterion (AIC) [Akaike, 1973], the C-index [Harrell et al., 1982], the area under the Receiver Operating Characteristic (ROC) curve (AUC) and the Integrated Brier Score (IBS) [Graf et al., 1999]. For AUC, six estimation procedures were implemented following the cumulative/dynamic (C/D) approach or the incident/dynamic (I/D) strategy: *survivalROC* (C/D) [Heagerty et al., 2000], *cenROC* (C/D) [Beyene and Ghouch, 2020], *nsROC* (C/D) [Pérez-Fernández et al., 2018], *smoothROCtime* (C/D and I/D) [Díaz-Coto et al., 2020], and *risksetROC* (I/D) [Heagerty and Zheng, 2005]. As AUC can be computed for each time point, Coxmos provides both AUC measurements across time and the average of AUC values for a set of time points.

The package also generates plots and performs statistical tests (t-test, ANOVA, Wilcoxon rank-sum test, and Kruskal-Wallis test) to evaluate and compare different models, identifying the best-performing ones according to the chosen performance metric.

#### 2.2.4 Model validation

The accomplishment of the proportional hazards (PH) assumption in Cox models can be assessed by analysing the Schoenfeld residuals. Coxmos provides residual plots and computes Schoenfeld’s statistical test by using functions from *survival* [Therneau, 2023] and *survminer* [Kassambara et al., 2024] R packages.

#### 2.2.5 Model interpretation

The Coxmos package incorporates several graphical tools for interpreting both Cox and PLS-based survival models, next summarized.

##### Forest Plot

Forest plots in Coxmos were adapted from the *survminer* R package [Kassambara et al., 2021], and summarize the estimated effects of covariates or PLS components in the final Cox models.

##### Predicted value distribution

Coxmos provides different types of predictions from the training data using the *survival* R package [Therneau, 2023]: linear predictor (lp) values, risk, expected number of events, and survival probability. Additionally, the distributions of predicted values grouped by event status are displayed in either density plots or histograms, allowing for an assessment of the model’s discriminative ability.

##### Contribution of predictive variables or components

The contribution to AUC of each original variable, PLS component, or linear predictor is plotted over a range of time points, to identify the most relevant variables for model’s performance.

##### sPLS visualization

PLS-based models can be interpreted through loading plots for variable contribution to components and relationships among variables, score plots with projected observations, or biplots, which combine variables and observations.

##### Pseudobeta coefficients

The estimated vector of Cox regression coefficients 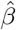 (Equation 1) reflects both the magnitude and direction of each predictor’s effect, i.e., whether they act as protective or risk factors. However, in sPLS-Cox models, the predictors included in the final Cox model are PLS components, which may hamper the interpretability of the effects of original predictor variables. To overcome this limitation, we defined the *pseudobeta coefficients* 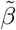 by leveraging the relationship between Cox regression coefficients 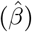 of the PLS components and the transformed weights (*W*^*^) of the original variables in the PLS model, which connect response and predictors in PLS models (Equation 9).

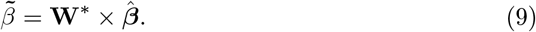

Coxmos tools for exploring pseudobeta coefficients include options to filter out non-significant components (according to a predefined significance threshold α), exclude variables with zero pseudobeta values, or highlight the top-contributing features. The percentage contribution of each variable to the final linear predictor can also be computed, accounting for cases in which a variable appears in multiple sPLS components. The total effect is then computed as the sum of its contributions across all components.

##### Kaplan-Meier curves

We adapted the Kaplan-Meier (KM) curves in the *survminer* R package [Kassambara et al., 2024] for all types of models and to show lp, PLS components or original variables. We combined this plot with a log-rank test. For numerical variables, Coxmos estimates the optimal cutoff to divide patients into lower and higher survival rates, and this cutoff can be used on test data to assess the model’s ability to differentiate potential risk groups.

#### 2.2.6 Survival prediction for new observations

Given a new set of patients with their corresponding covariate values, making predictions from a fitted Cox regression model is straightforward. However, for PLS-based survival models, the covariate values must be properly scaled and transformed into scores prior to obtaining such predictions. Thus, the score matrix **T**_pred_ for new observations with a covariate matrix **X**_test_ is computed as in Equation 10.

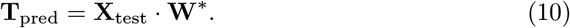

Most of the Coxmos functionalities mentioned in previous sections can also be applied to these predictions for new observations. For instance, the predicted value of a single new observation can be added to the training predicted value distribution to visualize its relative position. Additionally, the new patient’s pseudobeta coefficients can be compared with those from the trained model. In the case of multiple new observations (e.g. test data), the variable or component contribution to the model’s performance can be obtained to assess which variables have more impact on survival and to detect possible overfitting. As with single observations, pseudobeta coefficient plots can also be generated simultaneously for multiple patients. Finally, to evaluate model performance on unseen data, cutoff values from the training KM curves can be applied to stratify new patients and assess the model’s ability to separate them into groups with different survival probabilities, given by the log-rank test.

### 2.3 Model benchmarking

We compared the predictive performance of Coxmos survival models to machine learning (ML) methods on different datasets by using the C-Index and AUC metrics. An Analysis of variance (ANOVA) was applied to identify significant differences between methods and datasets (block factor), and least significant difference (LSD) intervals were used to identify the best-performing methods. Next we describe the models compared and how they were optimized.

#### 2.3.1 Coxmos models

All Coxmos models were optimized via 5-fold cross-validation with 3 repetitions on 80% of each dataset (training set). The performance metrics were computed on the remaining 20% observations (test set). The Cox model and CoxSW did not require hyperparameter tuning, whereas the two regularization parameters were optimized in the CoxEN model. For PLS-based methods, the tuned hyperparameters depended on the model: for sPLS-ICOX, both the number of PLS components and the penalty factor for variable selection; for sPLS-DRCOX and sPLS-DACOX, the number of PLS components and the number of variables to retain per component; and for (i)SB-sPLS and MB-sPLS models, the same as for their HD counterparts. The block-correlation matrix in MB-sPLS models was computed automatically by Coxmos.

The metric used for model optimization was the average AUC at 15 equally spaced time points. Penalty parameters were explored over the range 0.1-0.9 in increments of 0.2. For the number of variables to retain per component in the heuristic search, the minimum number of variables was set to 1, the maximum to the total number of variables, and the initial number of cutpoints to 10. The maximum number of PLS components considered was 8.

HD methods in Coxmos were also applied to MB datasets as an alternative strategy, after concatenating data blocks and applying soft-block scaling to each block *X*_*b*_ with *p*_*b*_ variables [Stevens et al., 2005] (Equation 11).

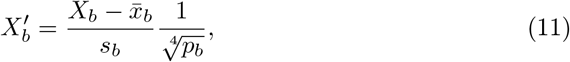

#### 2.3.2 Machine Learning models for survival analysis

To ensure a fair comparison between Coxmos and ML models, we again applied a 5-fold cross-validation with the same training and test partitions used for Coxmos models.

For HD datasets, we compared the following ML algorithms:

- **CoxBoost** [Binder and Schumacher, 2008], from the *CoxBoost* R package v1.5 [Binder, 2020]. We optimized the penalty value *λ*, whose initial value was set to the default, and the number of boosting steps, which ranged from 10 to 1000, with a maximum of 150 iterations.
- **MBoost** [Hofner et al., 2014], from the *mboost* R package v2.9.11 [Hothorn et al., 2024]. The number of boosting iterations was optimized, with a maximum of 1500 iterations.
- **Random Survival Forest (RSF)** [Ishwaran et al., 2008], from the *randomForestSRC* R package v3.4.1 [Ishwaran et al., 2023]. Three hyperparameters were optimized: the number of variables randomly selected at each split (mtry), the minimum size of terminal nodes (nodesize), and the number of trees (ntree). Since no built-in optimization function was available RSF, a two-step grid search strategy was implemented. First, a grid of values was defined for mtry (based on the square root of the number of predictors: 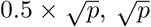 and 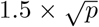) and nodesize (10, 15, 20). For each combination, a two-step search was performed to optimize the number of trees: an initial evaluation over four candidate values (250, 500, 750, and 1000), followed by a refinement step that evaluated ±50 trees around the best result. The combination of parameters with the highest average C-index across folds was selected. As no variable selection method is implemented in *randomForestSRC*, we also fitted an RSF model (RFS SV) on the variables with the highest importance, where the number of retained variables was set to the maximum number of variables selected by Coxmos models for the same dataset. For MB data, we considered the following ML methods:
- **IPF-LASSO** [Boulesteix et al., 2017], from the *ipflasso* R package v1.1 [Boulesteix et al., 2019]. Two versions were applied: *IPF LASSO*, with a fixed penalty factor of 1 per block; and *Penalized IPF LASSO*, where the penalty factor was optimized using the *cvr2*.*ipflasso* function in the package over a predefined range of values (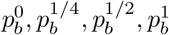, being *p*_*b*_ is the number of variables in each block). The penalty parameter *λ* was also optimized in both cases.
- **Priority LASSO** [Klau et al., 2018], from the *prioritylasso* R package v0.3.1 [Klau et al., 2023]. The penalty parameter *λ* was optimized internally, while the block priority order was optimized by evaluating all possible permutations.
- **Block Forest** [Hornung and Wright, 2019], from the *blockForest* R package v0.2.6 [Hornung and Wright, 2023]. Similarly to RSF, we applied a two-step grid search to determine the optimal number of trees. First, four predefined values (250, 500, 750, 1000) were evaluated, followed by a refinement step that evaluated ±50 trees around the best result. The number of trees that achieved the highest average C-index across folds was selected. This optimized value was used to train the final model with 300 iterations, allowing internal tuning of block-specific parameters using the *blockfor* function from the package. We also fitted a Block Forest model (Block Forest SV) on the variables with the highest importance, and the number of variables to retain was set to the maximum number of variables selected by Coxmos models for the same dataset.

### 2.4 Data

We describe next the datasets selected (summarized in Table 1) to test the HD and MB methods implemented in Coxmos and to compare them to ML models.

**Table 1:** Multi-omic studies analyzed in this work, including data modalities with their number of patients and variables, number of events, and EPV value.

| Dataset | Data Type | N. Patients | N. Var. | N. Event | EPV |
| --- | --- | --- | --- | --- | --- |
| BRCA | RNA-seq | 571 | 19318 | 68 | 0.004 |
| BRCA | miRNAs | 571 | 642 | 68 | 0.106 |
| BRCA | Proteomics | 571 | 518 | 68 | 0.184 |
| BRCA | Methylation | 571 | 17436 | 68 | 0.004 |
| BRCA | Clinical | 571 | 22 | 68 | 6.800 |
| HNSC | RNA-seq | 443 | 21520 | 152 | 0.007 |
| HNSC | miRNAs | 443 | 793 | 152 | 0.192 |
| HNSC | Mutations | 443 | 106 | 152 | 1.430 |
| HNSC | CNV | 443 | 57964 | 152 | 0.003 |
| HNSC | Clinical | 443 | 7 | 152 | 21.714 |
| HGSOC | RNA-seq | 290 | 12370 | 176 | 0.014 |
| HGSOC | miRNAs | 290 | 523 | 176 | 0.337 |
| HGSOC | CNV | 290 | 15055 | 176 | 0.012 |
| HGSOC | Methylation | 290 | 17441 | 176 | 0.010 |

1. **BRCA**: Multi-omic dataset from the TCGA repository [The Cancer Genome Atlas Network, 2012], which was preprocessed by our team and available at https://github.com/pilargmarch/multiomics2.0. It includes clinical information, RNA-seq expression data, miRNA-seq expression data, DNA methylation profiles, and protein levels for 571 patients with breast cancer. The preprocessing steps included removing variables with missing values, duplicates, and non-tumor samples. Additionally, patients with ambiguous clinical attributes, such as “Stage X” in AJCC pathologic staging, “not reported” prior malignancy status, or unspecified pathologic T or N stages, were excluded to ensure data quality and reliability.
2. **HNSC**: Multi-omic TCGA dataset [The Cancer Genome Atlas Network, 2015] with 443 Head and Neck Squamous Cell Carcinoma (HNSC) patients, which includes clinical data, RNA-seq gene expression, miRNA expression, mutations, and Copy Number Variations (CNV). The preprocessed datasets were obtained from [Bommert et al., 2022].
3. **HGSOC**: High Grade Serous Ovarian Cancer (HGSOC) multi-omic TCGA dataset [The Cancer Genome Atlas Network, 2011] for 290 patients containing RNA-seq gene expression, miRNA expression, CNVs, and methylation profiles. This dataset was downloaded and processed by our group as described in [Aguerralde-Martin et al., 2025].

Each dataset was prepared for survival analysis using our ATHAR R pipeline (https://github.com/BiostatOmics/ATHAR), and variables with missing data or inconsistencies were removed. Besides, Coxmos implements some of the ATHAR pipeline strategies to filter variables such as:

1. **Qualitative variables:** A Near Zero Variability (NZV) filter using the *caret* R package [Kuhn, 2008] that removes variables with few distinct values compared to the sample size. A variable *x*_*j*_ is removed when the following two conditions are met simultaneously:
  a. **Frequency Ratio**: 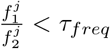, where 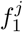 is the frequency of the most prevalent category for variable *x*_*j*_ and *f*_2_ is the frequency of the second most prevalent category. In our case, we set this threshold to *τ*_*freq*_ = 95*/*5.
  b. **UVP**: 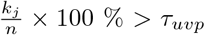, being *k* the number of classes of predictor *j* and *n* the sample size. We set the threshold to *τ*_*uvp*_ = 10 %.
2. **Quantitative variables:** Filter to remove near-constant variables with a Coefficient of Variation lower than *τ*_*CV*_. We set *τ*_*CV*_ = 0.1.

Finally, to illustrate Coxmos functionalities in more detail, we used the HGSOC study, which lasted 5481 days and where 60.69% of the 290 patients experienced the event (death).

## 3 Results and Discussion

### 3.1 Comparison of HD methods on clinical datasets

We first compared the performance of Cox and PLS-based models in Coxmos to ML methods on the available clinical datasets, where the full Cox regression model with no variable selection was also applicable. The ANOVA tests (Suppl. Tables S1 and S2) revealed a statistically significant effect of the dataset on model performance for both the C-Index (p-val < 0.0001) and AUC (p-val = 0.002) metrics, indicating that the characteristics of the data may influence the choice of the survival model. In contrast, almost statistically significant differences were found between methods for these metrics (C-Index p-val = 0.1087 and AUC p-val = 0.0802), suggesting a similar performance of the compared models on low HD data, such as the clinical data analyzed. Still, the consequently overlapping LSD intervals for the C-Index (Fig.2A) showed that PLS-based survival models (C-index mean 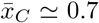) slightly outperformed the full Cox and CoxSW models 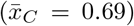 and the RF models 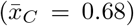, while CoxEN and ML boost models performed worse 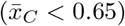. Among PLS-based models, sPLS-DRCOX achieved the highest average C-Index. For the AUC metric (Suppl. Fig. S1), we observed a superior average AUC for all Coxmos models 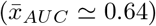 when compared to ML models 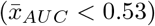.

**Figure 2:**
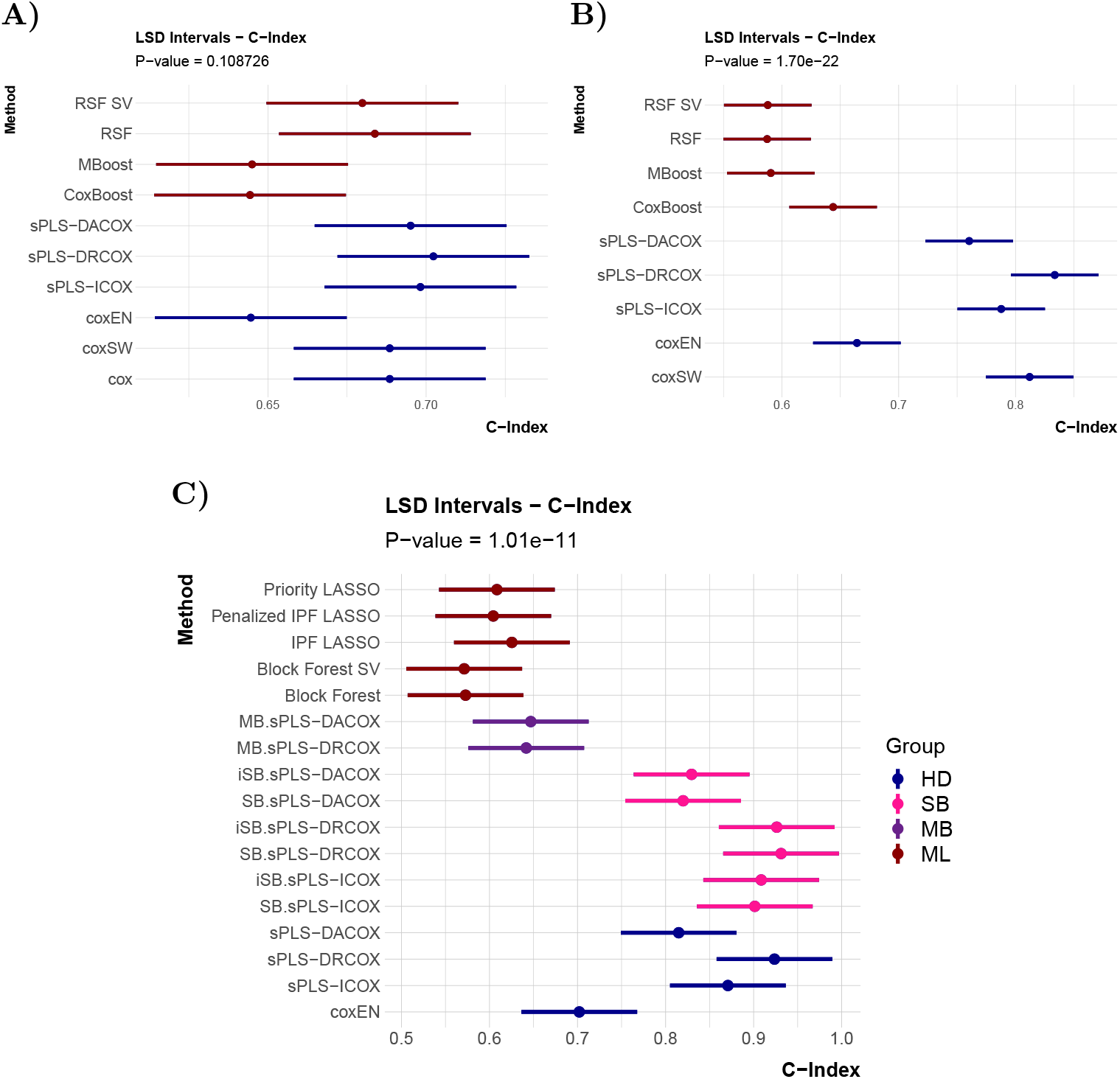
LSD intervals at 5% significance level for the C-Index, computed on test data. **A)** Comparison of HD and ML methods on clinical data from BRCA and HNSC studies. **B)** Comparison of HD and ML methods on omic data from BRCA, HNSC and HGSOC studies. **C)** Comparison of HD, MB and ML methods on multi-omic data from BRCA, HNSC and HGSOC studies.

### 3.2 Comparison of HD methods on single-omic datasets

The same HD survival models were compared on omic datasets, which present a much higher dimensionality than clinical data. Obviously, the full Cox regression model had to be excluded from this comparison.

Unlike the clinical data, the ANOVA test for omic data (Suppl. Tables S3 and S4) revealed statistically significant differences in model performance across both datasets and methods for the C-Index and AUC metrics (p-val *<* 0.0001). The C-Index LSD intervals (Fig. 2B) showed that all ML methods (with 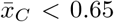) and coxEN 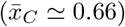 presented a significantly lower performance than the rest of the Coxmos methods (from 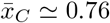 for sPLS-DACOX to nearly 0.84 for sPLS-DRCOX). AUC results (Suppl. Fig. S2) mostly confirmed these findings, but, in this case, coxEN performed similarly to its Coxmos counterparts. However, while sPLS-DRCOX obtained the highest average C-Index, sPLS-ICOX and coxSW achieved the highest AUC value 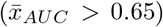. The overlapping LSD intervals for most Coxmos models, although indicating non-significant differences between them, suggest that Coxmos PLS-based survival models can properly handle extremely high dimensionality and multicollinearity present in omic data. Although our adapted version of coxSW yielded good results, it is worth noting that its application was not always feasible due to computational restrictions, as observed with CNVs in the HNSC dataset.

### 3.3 Comparison of MB methods on multi-omic datasets

Finally, we evaluated models on the multi-omic BRCA, HNSC and HGSOC studies. We considered four groups of models: HD methods in Coxmos applied to concatenated and block-scaled datasets; SB and MB models in Coxmos, which directly handle MB data; and ML models for MB datasets. Considering the size of these multi-omic datasets, full Cox and CoxSW models had to be excluded from the comparison. We also excluded non-MB ML methods due to their poor performance in previous comparisons.

The ANOVA tests (Suppl. Tables S5 and S6) revealed a significant effect of the dataset on the C-Index and AUC (p-val *<* 0.0001) as well as for the method (p-val *<* 0.0005). The C-Index LSD intervals (Fig.2C) showed that all PLS-based models in Coxmos, except MB.sPLS-DACOX and -DRCOX, outperformed ML models and coxEN. SB, iSB and concatenated sPLS-DRCOX and -ICOX obtained an average C-Index higher than 0.85, followed by SB, iSB and concatenated sPLS-DACOX 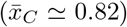, and coxEN 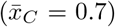. Coxmos MB and ML models 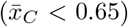 were significantly worse. For the AUC metric (Suppl. Fig. S3), the differences were less pronounced but ML-based methods still yielded the poorest performance, followed closely by MB approaches, while SB and HD models achieved the highest AUC scores. In particular, SB.sPLS-DRCOX and sPLS-DRCOX achieved the best AUC values 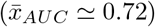.

Again, PLS-based survival methods in Coxmos showed their ability to efficiently analyse complex and HD data while properly addressing the multicollinearity issue. Interestingly, we found no clear superiority of MB methods over the concatenation strategy, or for the more flexible iSB models over their SB counterparts. The choice of the PLS-based strategy may depend on the characteristics of the dataset, highlighting the importance of having robust tools to optimize and compare different models, as those provided by Coxmos.

### 3.4 Case study on ovarian cancer

In this section, we showcase Coxmos’ functionalities by analysing the multi-omic HGSOC dataset. All figures and results shown were generated with the Coxmos R package.

An initial examination of the number of patients across time intervals and the proportion of them reaching the event of interest (Fig. 3) revealed a higher concentration of events at earlier time points. This may reduce statistical power and hinder reliable prediction at later time intervals, which have fewer observations.

**Figure 3:**
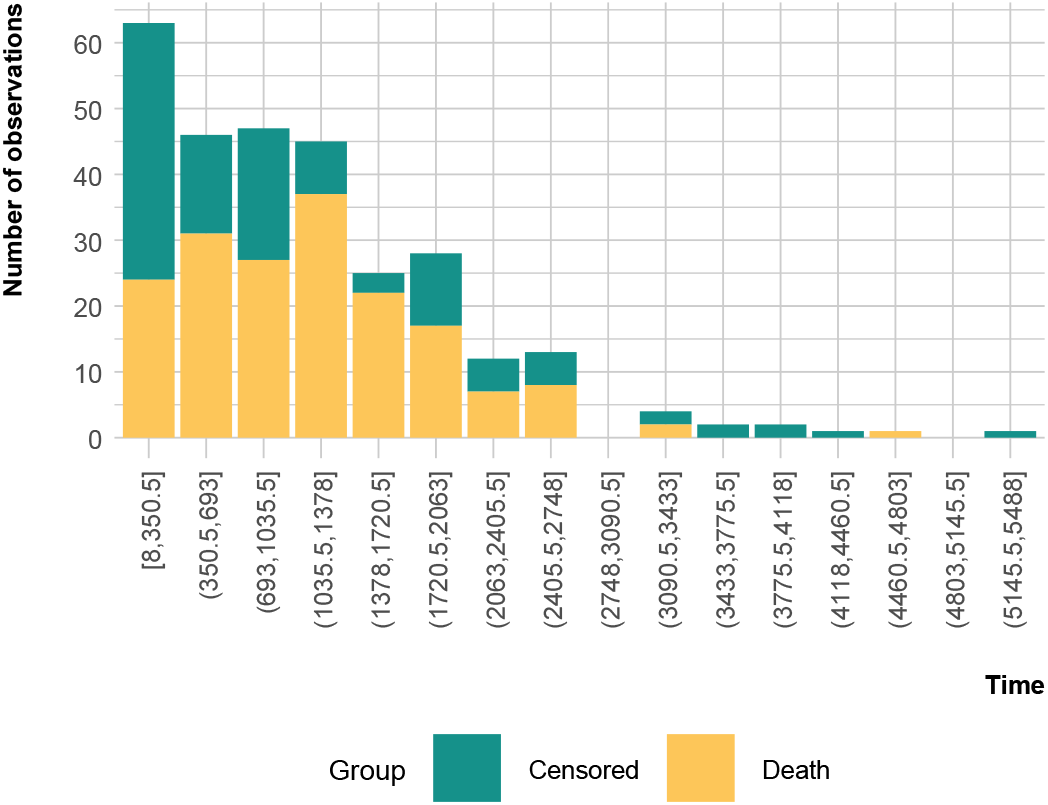
Distribution of patients and survival events per day interval in HGSOC dataset.

For model validation, the dataset was split into training (80%) and test sets (20%). The EPV values per omic block in the training set are below the commonly recommended threshold of 5–10 EPVs (Table 2), which reflects the pronounced high dimensionality of this dataset, requiring efficient multivariate approaches that can handle such HD data.

**Table 2:** EPV values per omic block for the training HGSOC dataset.

| Block | CNV | Genes | Methylation | miRNA |
| --- | --- | --- | --- | --- |
| EPV | 0.0093 | 0.0113 | 0.0080 | 0.2676 |

In this complex scenario, the optimization and evaluation of different survival models is essential, and Coxmos provides tools for facilitating this task. We compared the PLS-based survival methods in Coxmos –SB and MB methods, and HD methods on concatenated and scaled blocks– and coxEN. After hyperparameter tuning via 5-fold CV on training data with 3 repetitions, the C-Index performance metric was calculated for the optimized models on the test set (Fig. 4). The best predictive performance was achieved by sPLS-DRCOX (C-Index = 0.95), followed by SB.sPLS-DRCOX (C-Index = 0.92).

**Figure 4:**
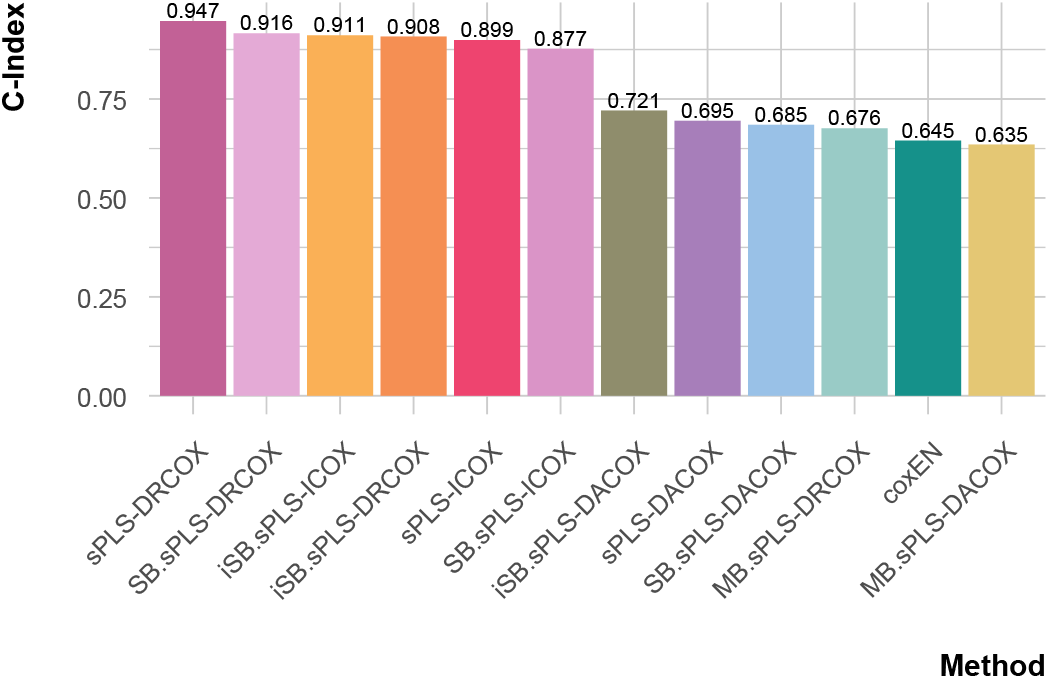
C-Index for survival models in Coxmos on the test set of the HGSOC dataset.

Model optimization yielded six statistically significant PLS components (Table 3) for the selected model (sPLS-DRCOX). Fig. 5A shows that the PH assumption is globally met (Schoenfeld’s test p-val = 0.1020), thus supporting its statistical validity for survival prediction. Although component 4 is statistically significant (p-val = 0.0053), the plot shows that Schoenfeld’s residuals are randomly distributed around zero without any clear temporal trend, suggesting no relevant PH assumption violation. All the significant components act as risk factors (Hazard Ratio *>* 1), as shown in Fig. 5B and Table 3. To assess each component’s predictive performance, Fig. 5C shows their contribution to the AUC in the training and test datasets. In the training set, the LP consistently achieved higher AUC values across time than any single component. In the test set, the first component outperforms the LP at earlier or later times, probably due to the lower number of observations at these time intervals, rather than a sign of overfitting. Despite this, the generally higher AUC of the LP suggests that multi-omic integration succeeds in capturing the biological signal. The discriminative ability of the model is also supported by the distributions of LP values for censored or event patients (Fig. 5D), as the event patients generally present higher LP values (lower survival probability) than the rest.

**Table 3:** Results for the sPLS-DRCOX model on HGSOC dataset.

| Variable | Coef | exp(Coef) | se(Coef) | Robust se | z | Pr(>\|z\|) |
| --- | --- | --- | --- | --- | --- | --- |
| comp_1 | 0.4977 | 1.6449 | 0.0409 | 0.0313 | 15.89 | $7.37 \times 10^{-57}$ |
| comp_2 | 1.3321 | 3.7892 | 0.1020 | 0.0892 | 14.94 | $1.89 \times 10^{-50}$ |
| comp_3 | 0.8525 | 2.3455 | 0.0733 | 0.0547 | 15.57 | $1.11 \times 10^{-54}$ |
| comp_4 | 0.5912 | 1.8062 | 0.0533 | 0.0481 | 12.30 | $9.21 \times 10^{-35}$ |
| comp_5 | 0.7498 | 2.1166 | 0.0670 | 0.0569 | 13.17 | $1.33 \times 10^{-39}$ |
| comp_6 | 0.7456 | 2.1077 | 0.0696 | 0.0707 | 10.55 | $5.27 \times 10^{-26}$ |

**Figure 5:**
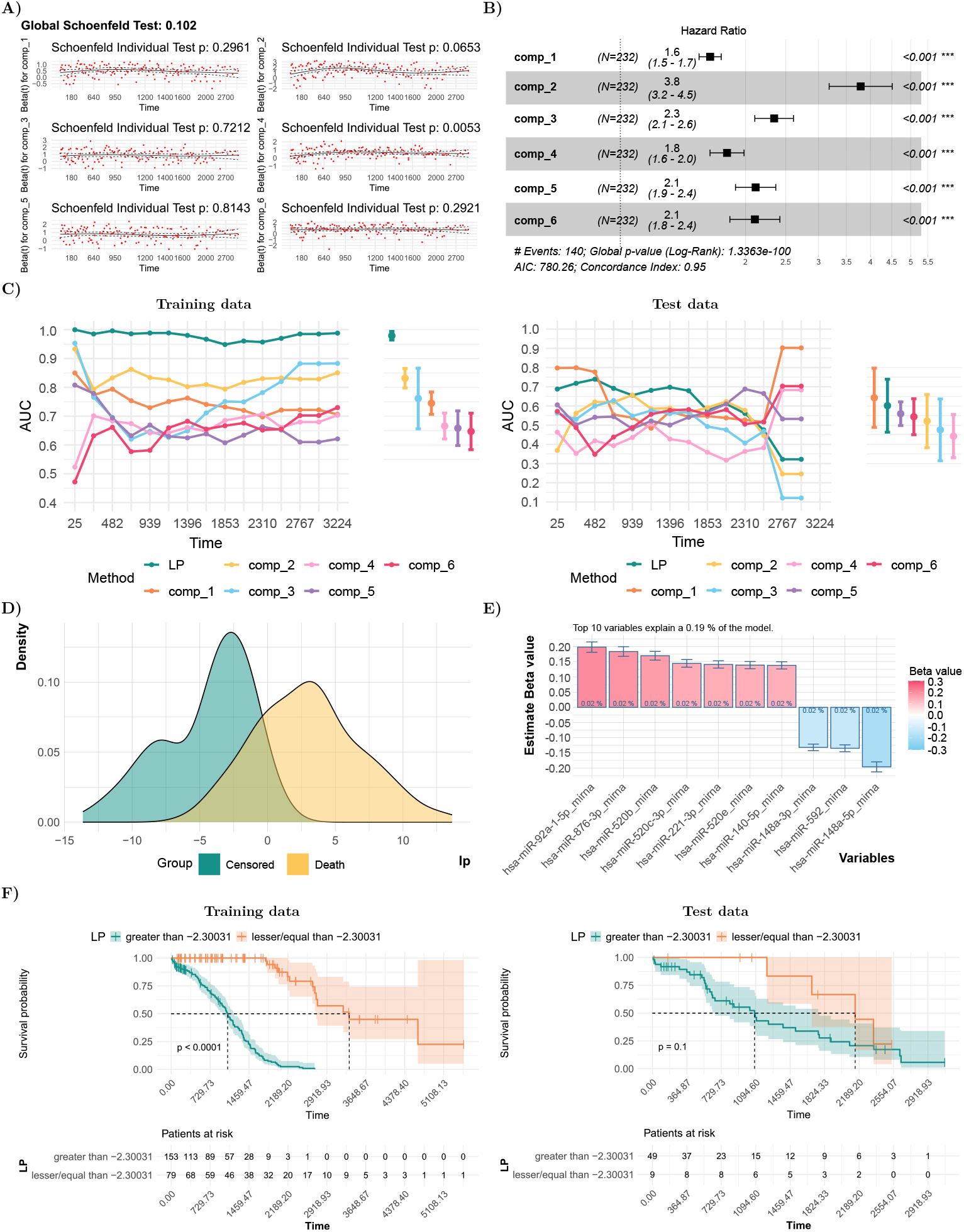
Coxmos results on the HGSOC dataset for the sPLS-DRCOX model. **A)** Schoenfeld residual test and plots for the model and significant PLS components. **B)** Forest plot showing the Hazards Ratio and 95% confidence intervals for the significant componentes. **C)** AUC decomposition into individual components using the training (left) and test (right) data. **D)** Density plots for the linear predictor (LP) per group. **E)** Estimated pseudobetas for the top 10 variables. **F)** Kaplan-Meier curve for the LP on the training (left) and test (right) data.

As PLS components are linear combination of the original values, in order to improve their interpretation, we developed a key feature of Coxmos: the pseudobeta coefficients for the original variables, which allow for identifying the most predictive variables in the model. Fig. 5E shows the top 10 variables with the highest pseudobetas, in absolute value. Interestingly, all of them are miRNAs, with seven of them acting as risk factors. [Resnick et al., 2009] reported higher serum levels of miR-92 in ovarian cancer patients versus controls, suggesting that miR-92 expression might be associated with poorer survival. The protective effect of miR-148a in the pseudobetas is supported by [Zhu et al., 2019, Zhao et al., 2016], who stated that overexpression of miR-148a reduces cell viability, migration, and invasion of ovarian cancer.

Finally, KM curves (Fig. 5F) showed that the LP clearly separates patients into low- and high-risk groups in the training set (log-rank p-val *<* 0.0001), while the separation is not so clear in the test set (p-val = 0.1) but maintains the direction of the effect. Notably, KM curves for the test set converge at late time points, where the number of observations declines, reducing precision and power, which may contribute to the non-significant result. These results highlight the usefulness of Coxmos’ functionalities for conducting a rigorous survival analysis, enabling the identification of both the strengths and weaknesses of the models.

## 4 Conclusion

In this work, we introduced Coxmos, a CRAN R package that provides a unified framework for survival analysis in HD and MB settings. Coxmos methods include the Cox Proportional Hazards model with stepwise (CoxSW) and ElasticNet (CoxEN) variable selection and,importantly, optimized PLS-based approaches specifically designed for HD and MB data. In addition, Coxmos incorporates useful tools for model optimization, validation, comparison, interpretation, and visualization to support reliable analyses and facilitate biological interpretation.

The comparison of methods across clinical, omic, and multi-omic datasets revealed several key insights. First, model performance in low-dimensional data, such as in clinical datasets, was more strongly influenced by the dataset itself than by the modeling strategy. Second, results on HD data, such as omic datasets, confirmed the advantages of our PLS-based strategies over other approaches, including ML methods. Third, MB methods showed a strong predictive potential and allow to better understand the contribution of each block to the model. Interestingly, concatenation with appropriate block scaling often achieved comparable results. MB PLS-based methods also generally outperformed state-of-the-art machine learning approaches. Combined with their enhanced interpretability, this makes them particularly suitable for survival analysis.

To demonstrate its practical application, Coxmos was applied to an ovarian cancer dataset, showcasing its ability to integrate multiple omic layers, identify relevant predictors, and assess their impact on patient survival. Moreover, Coxmos tools allow the detection of model limitations such as the lack of prediction generalizability due to reduced sample size or the need for careful validation.

In conclusion, Coxmos offers a robust and interpretable framework for survival analysis in complex data contexts, while highlighting the importance of model choice, optimization and validation.

## 5 Declarations

### Author contributions statement

S.T. and P.S. conceived the Coxmos idea. P.S. implemented the current version of Coxmos and performed the analyses. A.B.G. helped with the analyses. S.T. supervised the work. S.T., P.S. and A.B.G. wrote and reviewed the manuscript.

### Data availability

The databases used in this work are public and available at The Cancer Genome Atlas Program (TCGA) repository https://www.cancer.gov/ccg/research/genome-sequencing/tcga (2025).

### 5.1 Code availability

The Coxmos R package and two step-by-step tutorials for its application are freely available on the CRAN repository webpage at https://cran.r-project.org/web/packages/Coxmos/index.html and on GitHub at https://github.com/BiostatOmics/Coxmos. The code used for the analyses is available at https://github.com/BiostatOmics/Coxmos_scripts.

### 5.2 Ethics approval and consent to participate

Not applicable

### 5.3 Consent for publication

Not applicable

### 5.4 Materials availability

Not applicable

