## Supplementary information for "Coxmos: Interpretable survival models for high-dimensional and multi-omic data"

### Supplementary results

Pedro Salguero[1], Anabel Buendía-Galera[1], Sonia Tarazona[1,*]

[1]Department of Applied Statistics, Operations Research and Quality, Universitat Politècnica de València, Valencia, Spain.

### 1 Comparison of HD methods on clinical datasets

Table S1: ANOVA test results for clinical test data across BRCA and HNSC datasets and HD and ML methods for the C-Index metric. The factor Method refers to the applied survival method, and the factor Dataset represents the data used (block factor).

| | Df | Sum Sq | Mean Sq | F value | Pr(>F) | Significance |
|---|---|---|---|---|---|---|
| Dataset | 1 | 0.2191 | 0.2191 | 476.7481 | 0.0000 | *** |
| Method | 9 | 0.0098 | 0.0011 | 2.3575 | 0.1087 | |
| Residuals | 9 | 0.0041 | 0.0005 | | | |

Table S2: ANOVA test results for clinical test data across BRCA and HNSC datasets and HD and ML methods for the AUC metric. The factor Method refers to the applied survival method, and the factor Dataset represents the data used (block factor).

| | Df | Sum Sq | Mean Sq | F value | Pr(>F) | Significance |
|---|---|---|---|---|---|---|
| Dataset | 1 | 0.0796 | 0.0796 | 18.5352 | 0.0020 | ** |
| Method | 9 | 0.1029 | 0.0114 | 2.6649 | 0.0802 | . |
| Residuals | 9 | 0.0386 | 0.0043 | | | |

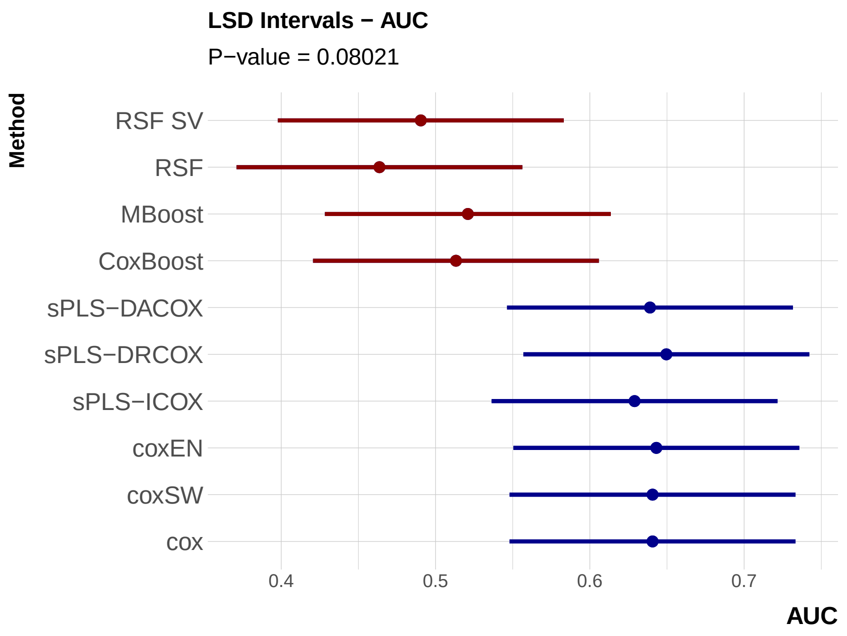


Figure S1: LSD intervals at 5% significance level for the AUC computed on clinical test data. Results for the compared HD and ML methods were obtained from clinical datasets in BRCA and HNSC studies.

### 2 Comparison of HD methods on single-omic datasets

Table S3: ANOVA test results for omic test data, across BRCA, HNSC, and HGSOC datasets and HD and ML methods, for the C-Index metric. The Method refers to the applied survival method, and the Dataset represents the data used.

| | Df | Sum Sq | Mean Sq | F value | Pr(>F) | Significance |
|---|---|---|---|---|---|---|
| Dataset | 11 | 0.4170 | 0.0379 | 9.9045 | 0.0000 | *** |
| Method | 8 | 0.9747 | 0.1218 | 31.8342 | 0.0000 | *** |
| Residuals | 85 | 0.3253 | 0.0038 | | | |

Table S4: ANOVA test results for omic test data, across BRCA, HNSC, and HGSOC datasets and HD and ML methods, for the AUC metric. The Method refers to the applied survival method, and the Dataset represents the data used.

| | Df | Sum Sq | Mean Sq | F value | Pr(>F) | Significance |
|---|---|---|---|---|---|---|
| Dataset | 11 | 0.4926 | 0.0448 | 12.0325 | 0.0000 | *** |
| Method | 8 | 0.4487 | 0.0561 | 15.0705 | 0.0000 | *** |
| Residuals | 85 | 0.3163 | 0.0037 | | | |

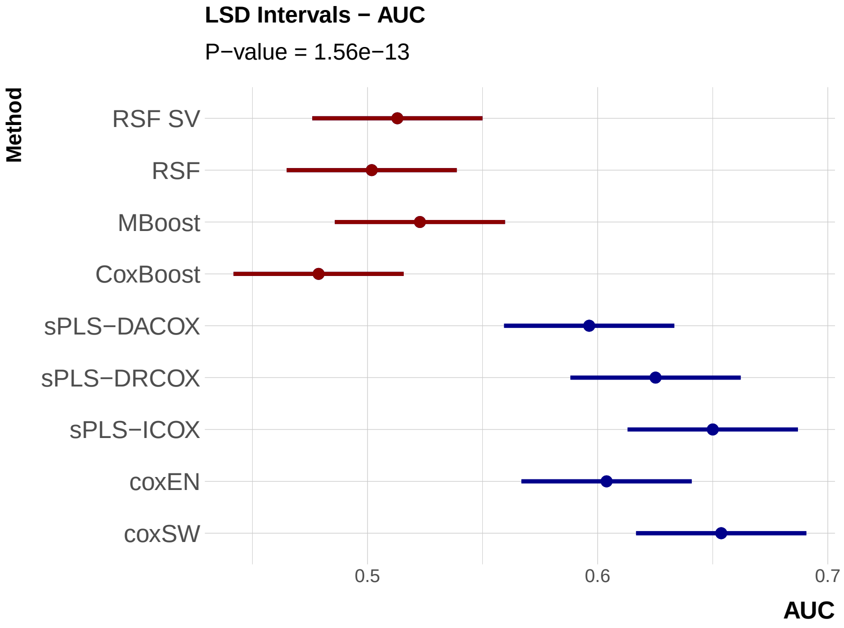


Figure S2: LSD intervals at 5% significance level for the AUC computed on single-omic test data. Results for the compared HD and ML methods were obtained from single-omic datasets in BRCA, HNSC and HGSOC studies.

### 3 Comparison of MB methods on multi-omic datasets

Table S5: ANOVA test results for multi-omic test data across BRCA, HNSC, and HGSOC datasets and HD, MB, and ML methods for the C-Index metric. The Method refers to the applied survival method, and the Dataset represents the data used.

| | Df | Sum Sq | Mean Sq | F value | Pr(>F) | Significance |
|---|---|---|---|---|---|---|
| Dataset | 2 | 0.1147 | 0.0573 | 17.6981 | 0.0000 | *** |
| Method | 16 | 0.9467 | 0.0592 | 18.2633 | 0.0000 | *** |
| Residuals | 32 | 0.1037 | 0.0032 | | | |

Table S6: ANOVA test results for multi-omic test data across BRCA, HNSC, and HGSOC datasets and HD, MB, and ML methods for the AUC metric. The Method refers to the applied survival method, and the Dataset represents the data used.

| | Df | Sum Sq | Mean Sq | F value | Pr(>F) | Significance |
|---|---|---|---|---|---|---|
| Dataset | 2 | 0.2788 | 0.1394 | 24.6774 | 0.0000 | *** |
| Method | 16 | 0.3569 | 0.0223 | 3.9495 | 0.0005 | *** |
| Residuals | 32 | 0.1807 | 0.0056 | | | |

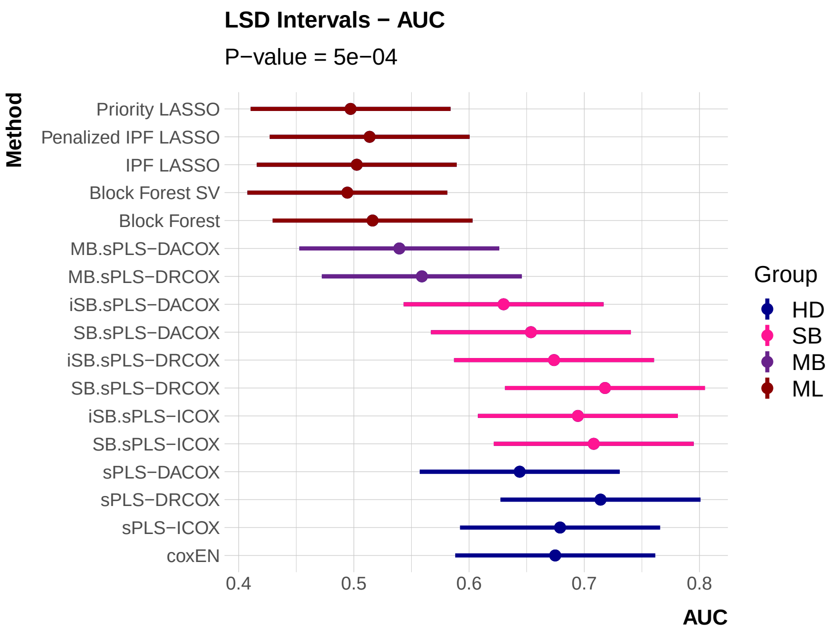


Figure S3: LSD intervals at 5% significance level for the AUC computed on multi-omic test data. Results for the compared HD, MB and ML methods were obtained from multi-omic datasets in BRCA, HNSC and HGSOC studies.
